# A Mathematical Model for Chemo-mechanically Induced Collective Cell Motility on Planar Elastic Substrates

**DOI:** 10.1101/2025.02.02.636116

**Authors:** Riham K. Ahmed, Tamer Abdalrahman, Neil H. Davies, Fred Vermolen, Thomas Franz

## Abstract

Cells interact with mechanical and chemical environmental cues, such as mechanical cues from other cells and chemical signals from growth factors. The current study aims to develop a mathematical model for combined chemically and mechanically induced collective cell motility on planar substrates. The mechanically induced cell motility is simulated using strain energy density gradients generated in an elastic substrate by cellular traction forces. For chemotaxis, Green’s function and Duhamel’s principle are used to solve the diffusion equation that describes the distribution of a growth factor and to represent chemo-mechanically induced deterministic collective cell motility on planar elastic substrates. Chemically induced motility of cells towards a growth factor source is predicted for different growth factor production and diffusion rates. Chemo-mechanical cues with varying growth factor production and diffusion rates are explored for the motility of four cells and one motile cell in the presence of one stationary cell. The developed model describes the chemo-mechanically induced motility of individual cells on planar substrates. The model provides valuable information for *in vivo* or *in vitro* studies due to its suitability for extension to other chemical source shapes, mobilised sources, many sources, and soluble concentration gradients.

## 1 Introduction

Cell motility is an essential feature of living cells [1]. Without cell motility, many biological processes cannot occur, including organ development and growth, wound healing, and the normal immune response to infection. Understanding mechanisms of cell motility is vital in the interpretation of emerging areas such as cellular transportation, tumour growth, the manufacture of artificial tissues in biotechnology, and regenerative medicine, which facilitate the in silico design of tissue engineering and regeneration [2–4]. Crawling movement is the most common mechanism cells utilise [5–7]. Collectively crawling cells respond to mechanical and chemical cues in their surrounding environment.

One process that influences collective cell motility is mechanical cell-cell communication (mechanotaxis). Cells in a collective environment interact mechanically with neighbouring cells in their environment and influence their motile behaviour. Cells adhere and apply mechanical traction forces to the substrate through the focal adhesions that deform the cellular environment. This traction force can be sensed by neighbouring cells [8], which can control the direction of motility according to various stimuli.

The motility of cells due to a chemical stimulus (chemotaxis) is a crucial mechanism by which tissues and organs become organised during development. Chemotaxis is involved in, for example, embryonic development, wound healing, and cancer metastasis. Promoting or inhibiting chemically induced cell motility can be desirable to support wound healing or prevent cancer progression.

Cells migrate towards or away from gradients of chemical sources, such as growth factors or bacterial agents, depending on the type of chemical signal [9]. Experimental studies illustrate that a gradient of a chemical signal orients cells [10–15]. It has been suggested that the movement of a cell in response to a chemical stimulus is driven by a ‘compass’, processing the signal in an intracellular mechanism that drives the cell in the direction of the gradient [16, 17]. However, other authors disregard this notion and propose that the cell orients itself by pseudopods [9].

Understanding the sensing mechanisms of cells is crucial for different biological processes, such as testing drug combination strategies and improving drug therapies. *In vivo* studies provide accurate descriptions of complex interactions and responses that occur within living organisms. However, ethical considerations, costs, resource constraints, and time complexity are challenges associated with these studies. *In vitro* experiments have become more sophisticated and complex and can generate accurate findings similar to *in vivo* studies. However, certain interactions or responses cannot be captured *in vitro*. Mathematical models and computational simulations can generate fast, reproducible, and controlled experiments at significantly lower costs and in line with 3R principles in animal research by predicting cell behaviours and guiding experimental design. Further, these models and simulations can provide valuable information and insights that are difficult to obtain experimentally. Combinations of computational modelling and *in vitro* experiments have opened new opportunities for researchers to directly obtain additional information on specific factors on individual and collective motility that can be evaluated only indirectly from *in vitro* experiments [18–21].

Mathematical models can be developed on different scales, ranging from population-based macroscopic to individual-based mesoscopic models. In this model, a microscopic cell-based model combining continuous and discrete approaches is developed. Continuous approaches are formulated in terms of partial differential equations (PDEs) to describe the motility of cell collectives as coarse-graining based on hydrodynamics principles [22–26]. These types of models help characterise the material properties of cell collectives using tissue-scale parameters where velocity, polarisation, and cell densities describe the evolution of cell motility. Spatially discrete approaches may involve discrete or continuous time steps. Discrete models consider individual cells in a cluster as particles with fixed or variable geometries during motility due to environmental cues, i.e., chemical, mechanical, or electrical cues [27–29]. These models efficiently track individual cells’ positions by computing adhesion, locomotion, repulsion, and drag forces between the centres of cells.

Mathematical and computational models that combine continuous and discrete approaches have been developed to study the effects of chemotactic cues on single [21, 30] and collective cell motility [31–35]. Most of these studies did not consider the multi-signalling stimuli, and the effects of mechanical cues on cell motility have been neglected. To combine the effects of chemotaxis and mechanotaxis, a mathematical description of the chemical stimulus requires the representation of the cell’s ability to sense chemical and mechanical cues and account for chemo-mechanical interactions.

The current study aims to develop a mathematical model for combined chemically and mechanically induced collective cell motility on planar substrates. The formulation uses an approach that considers individual cells in a colony, similar to particle simulations typical in biochemical problems, such as representing flow problems at the molecular scale.

## 2 Methods

The recently developed model for mechanically-induced stochastic cell motility [36] with **r***_i_*(*t*) = (*x_i_*(*t*), *y_i_*(*t*)) as the centre position of the cell *i* with radius *R* at a time *t* on the substrate, *ψ* is the total strain energy density, ẑ*_i_* is the normalised form of the motility unit vector, *ϱ*Δ***B***(*t*) is the Brownian motion and referred to as random walk where Δ***B***(*t*) ∼ *N*(0, Δ*t*), *ρ*(X) denotes the friction coefficient of the cell-substrate adhesion, and X denotes the position of a point in the cell that varies in the interval occupied by the cell, i.e., X = [−*R*, *R*],

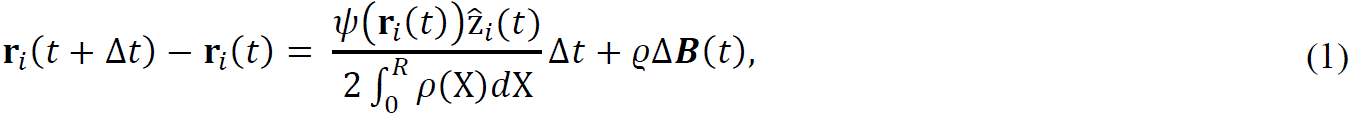

serves as a framework for chemo-mechanically induced cell motility by incorporating a diffusion PDE describing chemotaxis.

### 2.1 Chemically induced cell motility

The motility of cells on a 2D substrate in the presence of a growth factor gradient is investigated. The diffusion of a growth factor source (chemoattractant) that attracts the cells is approximated as a point source; hence, the point source is considered to be small compared to the cell areas. The chemoattractant point source is assumed to be stationary, and the diffusion field surrounding the point source is assumed to be instantaneous with time.

Suppose that **H** is the position of the point where the chemical source is released; then, the concentration of the growth factor that attracts the cells on the substrate is described by the following diffusion-reaction equation:

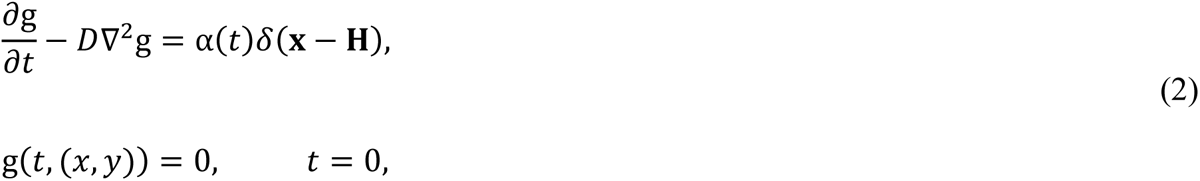

where g(*t*, (*x*, *y*)) denotes the concentration of the growth factor, α(*t*) is the amount of growth factor that is produced by the source per time, i.e., the production rate of the growth factor, and *D* is the diffusion coefficient of the growth factor. It is treated as a constant since the medium is assumed to be isotropic and homogeneous. Further, ∇^2^ = (*∂*^2^(⋅)/*∂x*^2^ + *∂*^2^(⋅)/*∂y*^2^) is the Laplace operator, and *δ*(*x, y*) represents the Dirac Delta distribution, which satisfies:

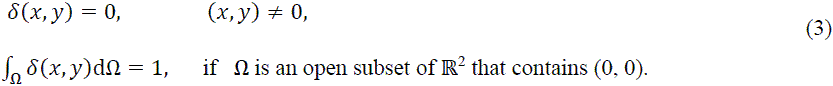

Initially, the growth factor concentration at all locations within the computational domain is assumed to be zero for simplicity. This assumption allows demonstrating the effect of the chemical gradient on cell motility as the release of the chemical gradient increases with time.

Green’s fundamental solution and Duhamel’s principle are used to solve Eq. (2) for the point source that triggers the release of the growth factor ([37], page 521). The fundamental solution in the 2D case for *t* > 0 is given by:

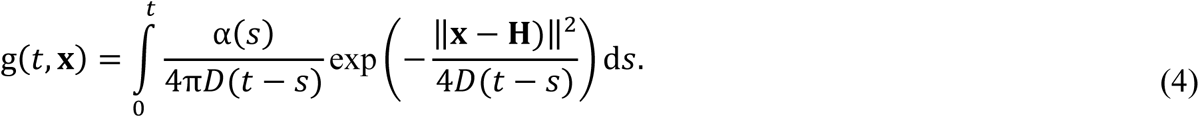

It is worth mentioning that the method of images can be used for a finite domain. The representation of Eq. (4) allows evaluation of the chemical concentration at any point on the substrate. However, it requires integration over an increasing time interval.

The diffusion of the growth factor results in a concentration gradient away from the site of the point source due to the particle movement from a region of high concentration to a region of low concentration. Therefore, cells sense the direction of the chemical gradient and migrate in the direction of the increasing gradient. Of course, it is also possible to use finite-element methods to solve this problem on a bounded domain. An advantage of using the fundamental solutions is that one can directly determine the concentration and its gradient at the position of the cell without the need to compute the concentration on the entire computation domain. Furthermore, the solution of Eq. (4) is not in H^1^(Ω) for a fixed time, which enlarges the error in the vicinity of the source. The concentration gradient of the chemical at any time *t* and position (*x, y*) is given by:

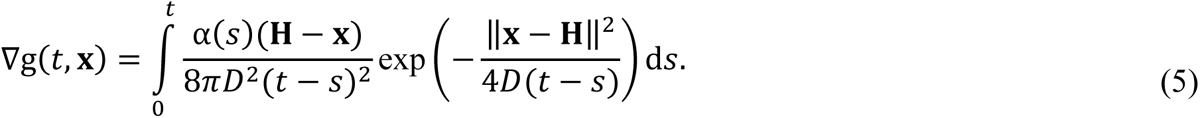

Since the gradient requires an integral over *s* from 0 to *t*, the actual concentration gradient of the growth factor is obtained by using the trapezoidal rule with the parameters listed in Table 1. For this purpose, the time is divided into a discrete set of points, *t_k_* = *k*Δ*t*, where Δ*t* is the time step.

**Table 1:** Parameters and values used in the model.

| Parameter | Description | Value and unit | Reference |
| --- | --- | --- | --- |
| $\alpha$ | Growth factor productivity (production rate) | 60 – 8,000 mol/$\mu\text{m}^2\text{s}$ | Estimated from Savinell [39] |
| $D$ | Growth factor diffusivity (diffusion rate) | 23 – 179 $\mu\text{m}^2/\text{s}$ | Estimated from Nauman [40] |
| $R$ | Cell radius | 50 $\mu\text{m}$ | [41] |
| $\sigma$ | Cell mobility in response to growth factor concentration | 1,000 $\mu\text{m}^4/\text{s mol}$ | Estimated from Vermolen [42], Vermolen and Gefen [43] |
| $\varrho$ | Probability of cell motion variability due to uncertainties | 0.4472 – 4.4721 | [36] |
| $\Delta t$ | Time step | 1 s | Estimated from Chen [44], Escribano [45], Peng and Vermolen [46], Chen [47] |

### 2.2 Chemo-mechanically induced cell motility

The growth factor point source is assumed to be located in a certain region of the substrate, and all cells are on one side of the growth factor source (Figure 1).

**Figure 1:**
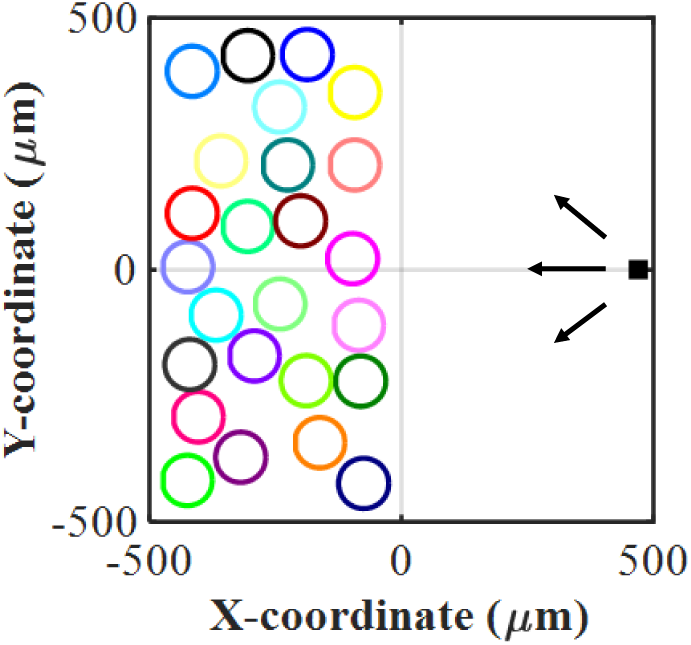
Initial positions of cells (open circles) and a growth factor point source (solid square). The growth factor diffuses through the 2D substrate and attracts the cells. The growth factor source is stationary and located at (470, 0), from where it diffuses in the direction of the cells.

For the combination of the two migratory mechanisms (mechanical and chemical), Eqs. (1) and (5) are combined to describe the influence of the growth factor stimulus on the mechanically induced cell motility. This gives the time-dependent displacement of the cells in the presence of the growth factor stimulus and random walk as:

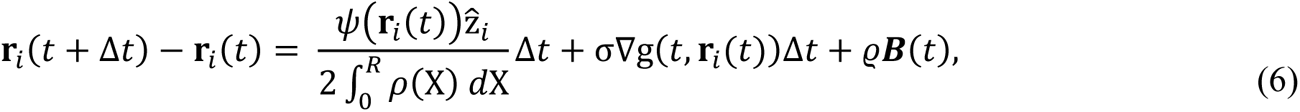

where σ denotes the mobility rate parameter of the cell’s response to the growth factor concentration gradient. This cell mobility rate σ can depend on spatial sensing over the cell body and its signalling rate. For simplicity, σ is assumed to be a constant in this model.

Each cell on the substrate is assumed to migrate due to sensing mechanical and chemical signals in Eq. (6). The reduction or obstruction of the chemotactic signal a cell receives from its neighbours is not considered in the current model, i.e., cells that are behind other cells still sense the same amount of chemotactic signal. The cells are modelled solely as sensing and responding to the growth factor gradient, without consuming, degrading, or secreting chemotactic signals at their spatial positions, such as T-cells. These cells primarily sense chemical gradients without substantially modifying the local concentration of the chemokines they detect [38].

The current model allows us to predict cell motility behaviour under different conditions and environmental properties by incorporating the effect of the chemical gradient in mechanically induced cell motility.

### 2.3 Cell motility simulations

The computational domain representing the planar substrate is 1,000 µm × 1,000 µm for all cell motility simulations. Open circles present the cells with a radius *R* = 50 µm, and the growth factor point source is represented by a solid square with a side length of 30 µm. The cells are initially located on the negative side of the X-axis of the substrate with dimensions 500 µm × 1,000 µm (Figure 1), where the chemical concentration is zero. Therefore, it is assumed that the release of the growth factor source that attracts the cells starts at *t* = 1 s. Further, it is assumed that the growth factor source is located at point (470, 0) µm and causes a point-symmetric chemical gradient over the 2D space. The dimensions of the computational domain are estimated from Giniūnaitė [48] and Mousavi [49].

The simulations mimic hypothetical cases; hence, hypothetical values have been used for many input parameters. The values of the model parameters introduced in the above equations are listed in Table 1 unless stated otherwise.

Each simulation experiment runs with a time step of 1 s. It is worth noting that the cells move around the chemical source randomly once they reach the chemical source (at X ≈ 470). Cells are allowed to have 30% overlap during the chemically induced motility of cells to account for cell deformation due to cell-cell contact [46]. Cell-cell contact corresponding to the strain energy density is considered in the current model to prevent 100% cell overlap during cellular mechanical interactions.

First, the evolution of the growth factor concentration gradient on a single is investigated for different values of the production rate (α) and diffusion rate (*D*), respectively. Second, the motility due to the chemical concentration gradient and the effect of the growth production rate and diffusion rate is demonstrated. Finally, the motility of cells as a response to the chemical concentration gradient and mechanical cellular interaction is studied.

The numerical results are obtained using MATLAB (The MathWorks, Inc., Natick, MA, USA).

### 2.4 Quantitative validation

To facilitate comparison with experimental data, the model outputs are scaled by adjusting the mobility parameter σ_cal_ to 2.76 µm^4^/s mol, such that the predicted mean velocities fall within reported experimental ranges (≈ 0.1-0.85 µm/min) for fibroblast migration on planar substrates [50, 51]. This calibration is performed using a representative reference condition corresponding to production rate α = 970 mol/µm^2^s and diffusion rate *D* = 179 µm^2^/s in the chemically induced collective cell motility (25 cells), for which the original model predicts a mean directed migration velocity (*υ*_model_) of approximately 54.26 µm/min. Taking a representative experimental value (*υ*_exp_) of 0.15 µm/min corresponding to the average velocities for fibroblast migration [50, 51]:

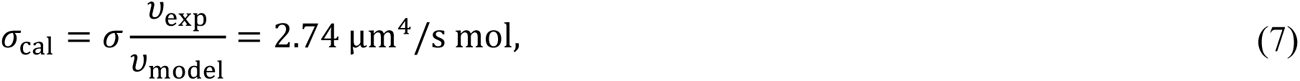

This calibration does not alter the qualitative behaviour of the model or the dependence of the cell motility on the growth factor production and diffusion rates. It rather ensures that the model operates within physiologically relevant regimes. Such scaling approaches are commonly used in computational models when the governing equations capture relative behaviour but lack intrinsic limits on absolute velocity.

## 3 Results

### 3.1 Chemically induced single and collective cell motility

#### 3.1.1 Effect of growth factor productivity and diffusivity on chemically induced motility velocity of a single cell

The migratory displacement of a single cell with fully deterministic motion towards a growth factor source on a substrate is considered for different growth factor production rates and diffusion rates. The time required for the cell to migrate to the growth factor source decreases with increasing growth factor productivity (Figure 2a) and diffusivity (Figure 2b). The initial distance between the cell and the growth factor source is fixed to 900 µm for each case. The final distance between the cell and the growth factor is smaller than the cell radius (*R* = 50 µm), i.e., the cell reaches the growth factor source since repulsive forces are not considered.

**Figure 2:**
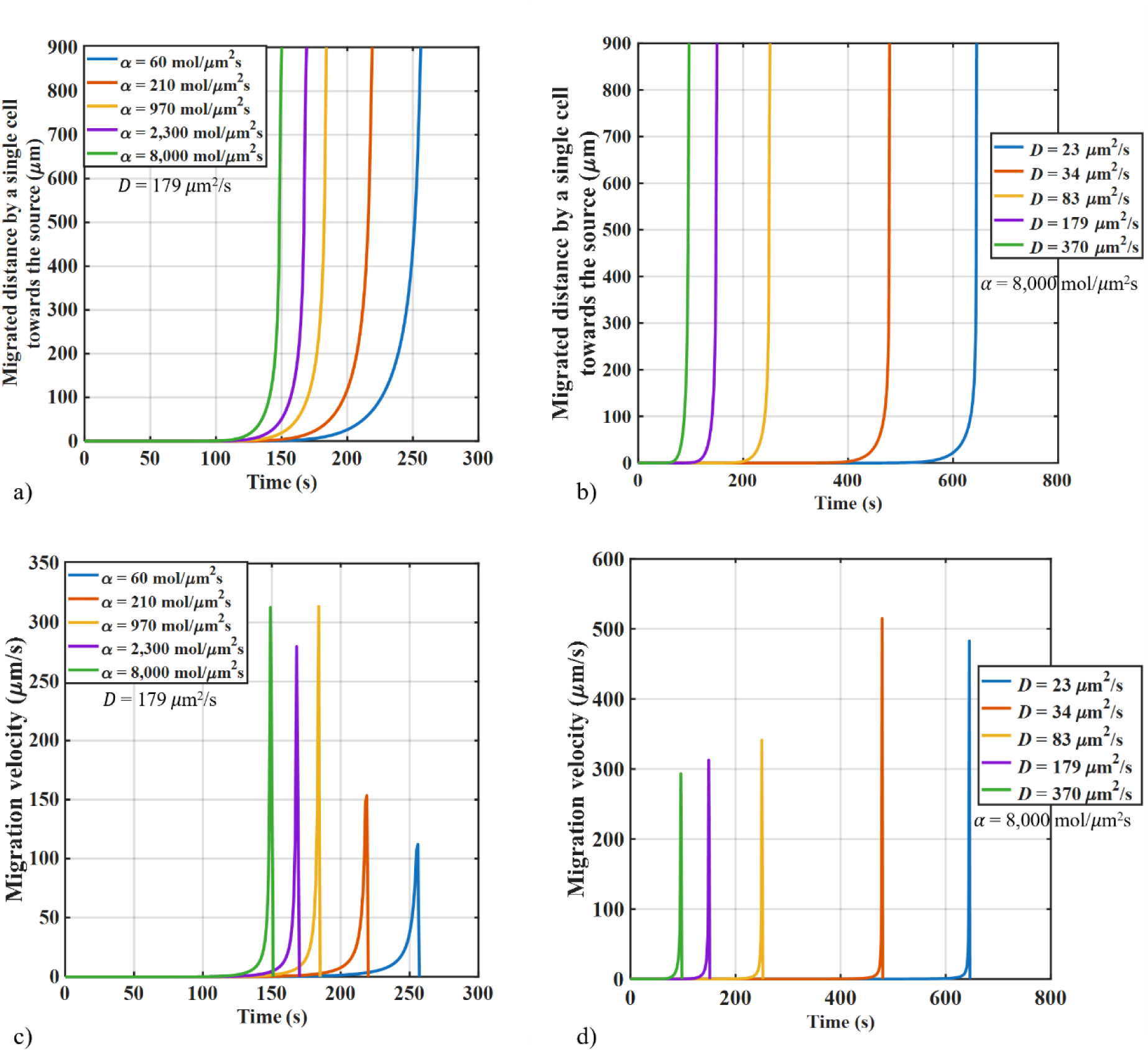
Chemically induced motility of a single cell. Distance between cell centre and growth factor source versus time of a cell centre reaching the growth factor source in fully deterministic motion (i.e., absence of stochastic motion) on a planar substrate for different values of the growth factor production rate (a) and diffusion rate (b). The time required for the cell to migrate to the growth factor source decreases with increasing growth factor production rate α (a, for *D* = 179 µm^2^/s) and growth factor diffusion rate *D* (b, for α = 8,000 mol/µm^2^s). Velocity versus time of a cell centre migrating towards the growth factor source in fully deterministic motion on a planar substrate for different values of (c) the growth factor production rate α with constant growth factor diffusion rate *D* = 179 µm^2^/s and (d) the diffusion rate *D* with constant α = 8,000 mol/µm^2^s (b). An increase in the growth factor production rate α causes an increase with subsequent plateauing of the maximum migration velocity of the cell, and shift of the peak velocity to earlier times (c). An increasing diffusion rate *D* results in an initial increase followed by a decrease in the maximum cell migration velocity and an earlier occurrence of the peak velocity (d). (Values and sources of the model parameters are provided in Table 1.)

The motility velocity of a single cell with fully deterministic motion towards the growth factor point source on a 2D substrate is determined for different growth factor production rates and diffusion rates. The results suggest that an increase in the growth factor production rate causes an increase with subsequent plateauing of the maximum migration velocity of the cell, and a shift of the peak velocity to earlier times (Figure 2c). An increasing diffusion rate results in an initial increase followed by a decrease in the maximum cell migration velocity, and an earlier occurrence of the peak velocity (Figure 2d).

#### 3.1.2 Chemically induced collective cell motility for varying growth factor productivity

Twenty-five cells are randomly located on a half-region of the substrate with a dimension of 1,000 µm × 1,000 µm, and a growth factor source with a constant diffusion rate of *D* = 179 µm^2^/s is used to study the effect of growth factor production rate on cell motility. Fully deterministic motion (i.e., without random movement) is considered. Three different growth factor production rates are considered: α = 60, 970, and 8,000 mol/µm^2^s.

For the growth factor source with α = 60 mol/µm^2^s (Figure 3a), the cells closest to the growth factor source start migrating at *t* = 70 s after the growth factor release is started. The two cells initially closest to the growth factor source reach the growth factor source first (*t* = 133 s). The cells with initial positions furthest from the growth factor source do not detect the chemical concentration gradient (*t* = 200 s). Once the cells have reached the growth factor source, they stay close to the source and keep moving around it along the Y-axis. All cells reach the growth factor source at *t* = 294 s. The cells in contact deform (shown as overlap) as inter-cellular forces are not considered in the model in combination with the chemical signal (see also supplemental video V3A).

**Figure 3:**
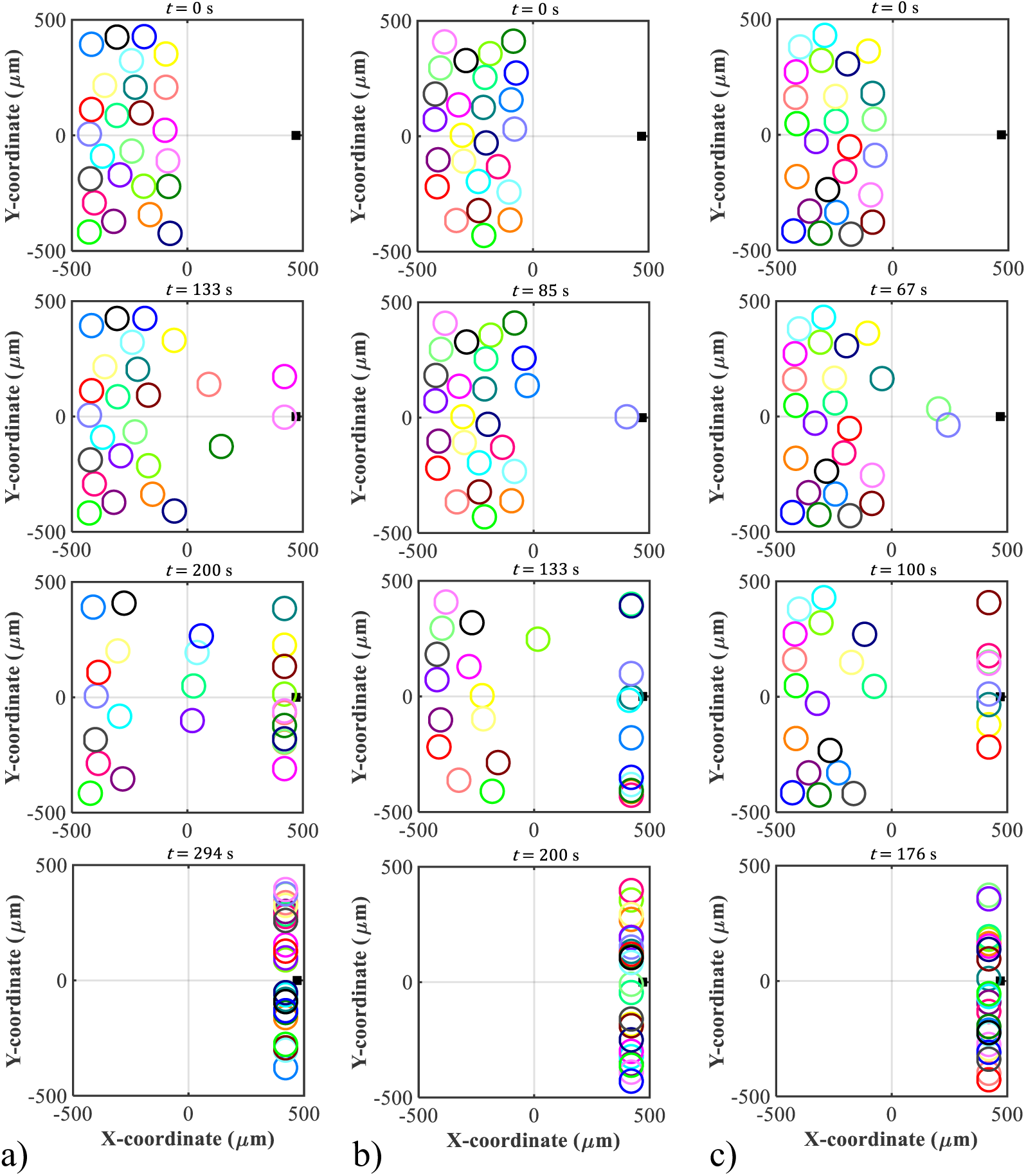
Chemically induced collective cell motility for varying growth factor productivity. Positions of 25 cells (open circles) migrating with fully deterministic motion on a 2D planar substrate towards a growth factor source (solid square) with a constant diffusion rate *D* = 179 µm^2^/s and different production rates of α = 60 mol/µm^2^s (a), α = 970 mol/µm^2^s (b), and α = 8,000 mol/µm^2^s (c). The growth factor source is located on the substrate region with X > 0 (right), and the initial arrangement of the cells on the substrate region with X < 0 (left) is the same for the three cases (at *t* = 0 s). a) For α = 60 mol/µm^2^s, the closest cells to the growth factor source start migrating towards the source, and two cells reach the source first at *t* = 133s. At *t* = 200 s, the cells furthest from the growth factor source do not migrate towards the growth factor source as they do not detect the chemical concentration gradient (also see supplemental video V3A). The cells overlap in the absence of repulsive cellular interactions. Once the cells reach the growth factor source, they move around it randomly along the Y-axis. At *t* = 294 s, all cells reach the growth factor source. b) For α = 970 mol/µm^2^s, one cell reaches the growth factor source at *t* = 58 s. At *t* = 133 s, the closest cells reach the growth factor and move around it randomly along the Y-axis. The cells furthest from the growth factor source do not migrate towards the source. The cells overlap in the absence of repulsive cellular interactions (see also supplemental video V3B). At *t* = 200 s, all cells reach the growth factor source. c) For α = 8,000 mol/µm^2^s, the closest two cells reach the growth factor source at *t* = 67 s. At *t* = 100 s, the closest cells reach the source and move around it randomly along the Y-axis. The cells furthest from the growth factor source do not migrate towards the source (see also supplemental video V3C). The cells overlap in the absence of repulsive cellular interactions. At *t* = 176 s, all cells reach the growth factor source. (Values and sources of the model parameters are provided in Table 1.)

The growth factor source with α = 970 and 8,000 mol/µm^2^s results in similar cell behaviour as the growth factor source with α = 60 mol/µm^2^s. In the final stage, all cells reach the growth factor source at *t* = 200 s for α = 970 mol/µm^2^s (Figure 3b) and at *t* = 176 s for α = 8,000 mol/µm^2^s (Figure 3c), demonstrating faster migration at higher production rates α. The cells start migrating earlier towards the growth factor source and reach faster motility velocities for a high production rate *α* compared to a lower production rate (see also supplemental videos V3B and V3C).

#### 3.1.3 Chemically induced collective cell motility for varying growth factor diffusivity

Collective cell motility due to a growth factor source on a planar substrate with three different growth factor diffusion rates of *D* = 23, 83, and 179 µm^2^/s and a constant growth factor production rate of α = 8,000 mol/µm^2^s is investigated using 25 cells arranged randomly in the region X < 0 of the substrate with the dimensions of 1,000 µm × 1,000 µm. The random movement of the cells is not considered.

For *D* = 23 µm^2^/s, cells start moving at *t* = 240 s after the growth factor release is started (Figure 4a). Only two cells move towards the growth factor source until the closest single cell reaches the growth factor source (*t* = 315 s). After that, the closest cells to the growth factor source reach the source and continue to move along the Y-axis (*t* = 485 s). The cells contact each other and deform (represented by overlap) without repulsive inter-cellular forces (see also supplemental video V4A). All cells reach the growth factor source at *t* = 836 s.

**Figure 4:**
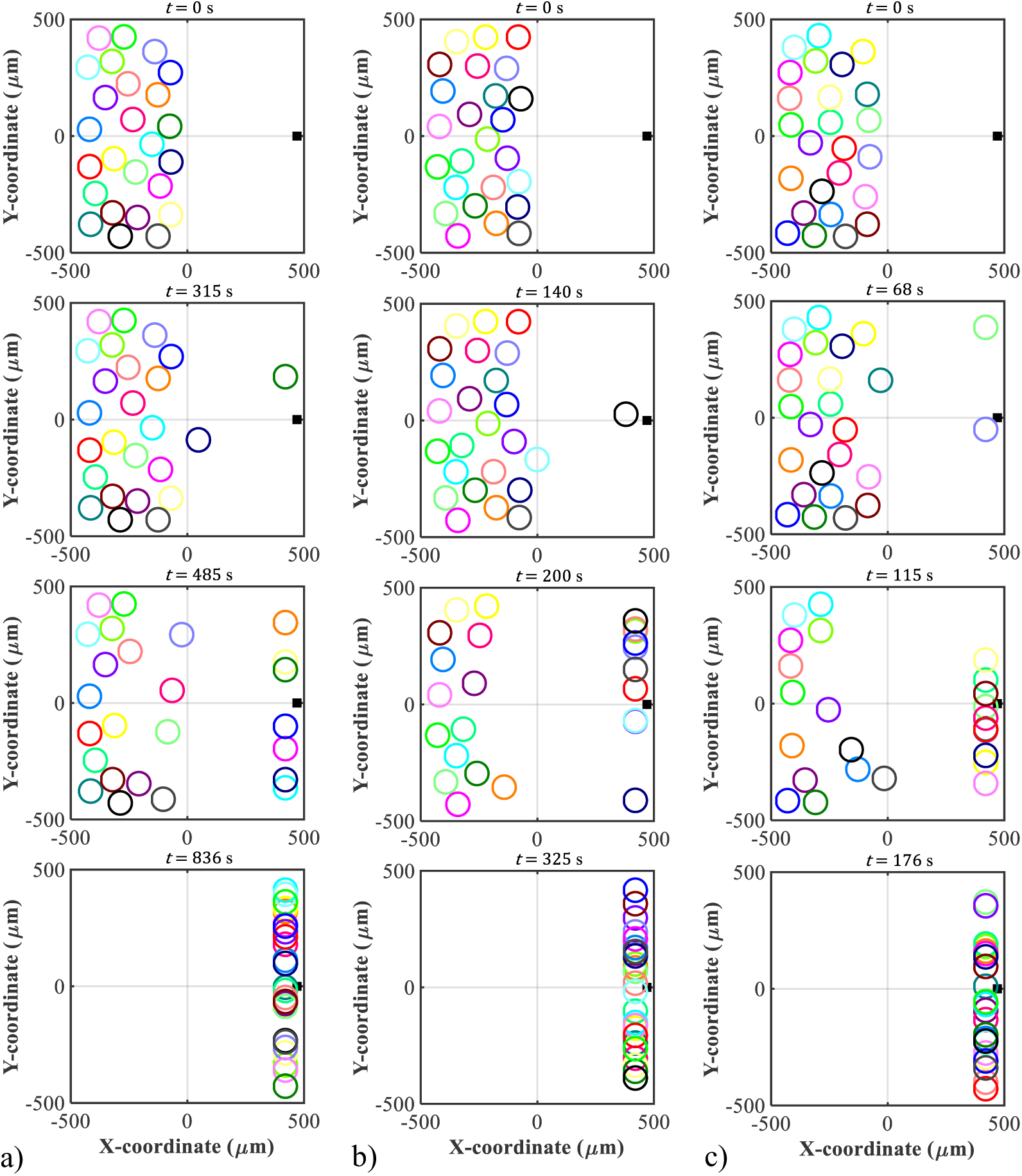
Chemically induced collective cell motility for varying growth factor diffusivity. Positions of 25 cells (open circles) migrating with fully deterministic motion on a 2D planar substrate towards a growth factor source (solid square) with a constant production rate α = 8,000 mol/µm^2^s and different diffusion rates of *D* = 23 µm^2^/s (a), *D* = 83 µm^2^/s (b), and *D* = 179 µm^2^/s (c). The growth factor source is located on the substrate region with X > 0 (right), and the initial arrangement of the cells on the substrate region with X < 0 (left) is the same for the three cases (at *t* = 0 s). a) For *D* = 23 µm^2^/s, the closest two cells to the growth factor source start migrating towards the source, and one cell reaches the growth factor source first at *t* = 315 s. At *t* = 485 s, the cells furthest from the growth factor source do not migrate towards the source as they do not detect the chemical concentration gradient (see also supplemental video V4A). Once the cells reach the growth factor source, they move around it randomly along the Y-axis. The cells overlap in the absence of repulsive cellular interactions. At *t* = 836 s, all cells reach the growth factor source. b) For *D* = 83 µm^2^/s, only two cells migrate towards the growth factor, and one cell reaches the source at *t* = 140 s. At *t* = 200 s, the closest cells reach the growth factor and continue moving around it randomly along the Y-axis (see also supplemental video V4B). The cells furthest from the growth factor source do not migrate towards the source. The cells overlap in the absence of repulsive cellular interactions. At *t* = 325 s, all cells reach the growth factor source. c) For *D* = 179 µm^2^/s, the closest cells start migrating towards the growth factor source, and two cells reach the source at *t* = 68 s. At *t* = 115 s, the cells furthest from the growth factor source do not migrate to the source (see also supplemental video V3C). The cells overlap in the absence of repulsive cellular interactions. After *t* = 176 s, all cells reach the source and move around it randomly along the Y-axis. (Values and sources of the model parameters are provided in Table 1.)

For *D* = 83 µm^2^/s (Figure 4b), only two cells initially move towards the growth factor source, and one cell reaches the source first at *t* = 140 s. The closest cells to the growth factor source reach the source and keep moving around it along the Y-axis (*t* = 200 s). The cells around the growth factor contact each other and deform (represented by overlap) without repulsive inter-cellular forces. Finally, all cells migrate and reach the growth factor source after *t* = 325 s (see also supplemental video V4B).

For *D* = 179 µm^2^/s (Figure 4c), the closest cells to the growth factor source migrate first towards the source, and two cells reach the source first (*t* = 68 s). The cells furthest from the growth factor source do not move as they do not detect the chemical concentration gradient (*t* = 115 s). Once the cells reach the growth factor source, they move around randomly along the Y-axis (see also supplemental video V4C). All cells reach the growth factor source after *t* ≈ 176 s, whereas this takes *t* ≈ 325 s for *D* = 83 µm^2^/s and *t* ≈ 836 s for *D* = 23 µm^2^/s, indicating faster migration at the higher growth factor diffusion rates considered in this simulation.

### 3.2 Chemo-mechanically induced collective cell motility

#### 3.2.1 Chemo-mechanically induced collective cell motility for varying growth factor productivity

Four cells migrating due to mechanical cues and a chemical signal are considered on a substrate with three different growth factor production rates of α = 60, 970, and 8,000 mol/µm^2^s and a constant diffusion rate of *D* = 179 µm^2^/s. Initially, the cells are located in a rhomboid arrangement in the substrate region X < 0, with an equal rhombus side length of 327 µm, a minor axis length of 260 µm, and a major axis length of 600 µm (Figure 5, *t* = 0 s).

**Figure 5:**
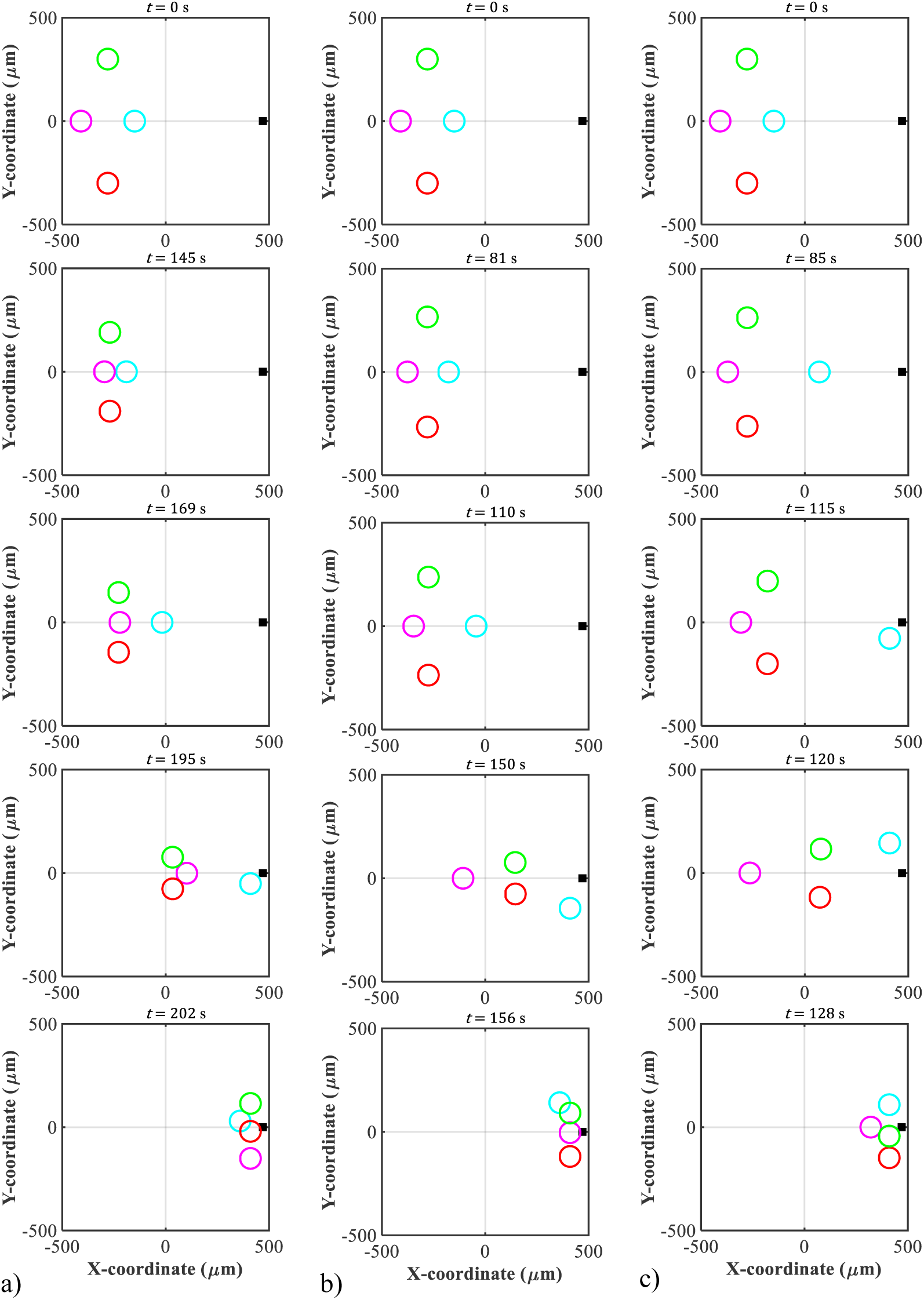
Chemo-mechanically induced collective cell motility for varying growth factor productivity. Positions of four cells (open circles) on a substrate during motility with no random motion towards a growth factor source (solid square) with constant diffusion rate *D* = 179 µm^2^/s and different production rates of (a) α = 60 mol/µm^2^s, (b) α = 970 mol/µm^2^s, (c) α = 8,000 mol/µm^2^s. The growth factor source is located on the substrate region with X > 0 (right). a) Initial positions of the cells at *t* = 0 s on the substrate region with X < 0 (left) in rhombus shape with a side length of 327 µm, minor axis of 260 µm, and major axis length of 600 µm. For α = 60 mol/µm^2^s, the cells move towards each other in the direction of the resultant strain energy density gradient at *t* = 145 s. Neighbouring cells contact each other and move towards the growth factor source (see also supplemental video V5A). At *t* = 169 s, front cell moves faster towards the growth factor source; hence, the chemical signal is more dominant than the mechanical cues. The front cell reaches the source, and other cells contact each other and cluster towards the growth factor source at *t* = 195 s. At *t* = 202 s, the cells reach the growth factor source and move around it along the Y-axis. The cells overlap in the absence of repulsive cellular interactions. b) Initial positions of the cells at *t* = 0 s for α = 970 mol/µm^2^s. At *t* = 81 s, neighbouring cells move towards each other following the resultant strain energy density gradient. At *t* = 110 s, the front cell moves towards the growth factor source, and other cells continue moving towards each other in the direction of the source. The front cell reaches the growth factor source, and the closest cells to the source move faster towards the source at *t* = 150 s (see also supplemental video V5B). At *t* = 156 s, the cells reach the growth factor source. The cells overlap in the absence of repulsive cellular interactions. c) Initial positions of the cells for α = 8,000 mol/µm^2^s. At *t* = 85 s, the cells move towards the chemical concentration gradient (see also supplemental video V5C). At *t* = 115 s, the front cell reaches the growth factor source first. The closest cells to the source move faster towards the source at *t* = 120 s. At *t* ≈ 128 s, the cells reach the growth factor source. (Values and sources of the model parameters are provided in Table 1.)

For α = 60 mol/µm^2^s (Figure 5a), the cells primarily move towards each other in the dominant direction of the resultant strain energy density gradient in the substrate until two cells are in contact at Y = 0 (*t* = 145 s). After that, the cell nearest to the growth factor source starts moving towards the source, and the other three cells continue moving towards each other and the growth factor source (*t* = 169 s). After the first cell reaches the growth factor source, the three other cells aggregate and move collectively towards the source until the chemical signal is more dominant than the mechanical cues (*t* = 195 s). Finally, the cells reach the growth factor source and impinge on each other since repulsive forces are not considered (*t* ≈ 202 s) (see also supplemental video V5A).

For α = 970 mol/µm^2^s, the cells move towards each other initially after *t* = 81 s (Figure 5b). The cells continue moving towards the growth factor source (see also supplemental video V5B); hence, the chemical signal is more dominant than the cellular mechanical interaction at *t* = 110 s. The cells closest to the growth factor source move faster and reach the source first after *t* = 150 s. Finally, all four cells reach and cluster around the growth factor source after *t* = 156.

For α = 8000 mol/µm^2^s (Figure 5c), the cells move slightly towards each other (see also supplemental video V5C) until the chemical signal is more dominant than the mechanically induced motility of cells (*t* = 85 s). The front cell reaches the growth factor source first (*t* = 115 s) and the cells closer to the growth factor source move faster towards the source than the more distant cell (*t* = 120 s). The cells move randomly around the source along the Y-axis. The distant cell reaches the source last, and the cells cluster around the source (*t* ≈ 128 s).

#### 3.2.2 Chemo-mechanically induced collective cell motility for varying growth factor diffusivity

Four cells are considered on a substrate with three different growth factor diffusion rates, i.e., *D* = 23, 83, and 179 µm^2^/s, and a constant production rate of α = 8,000 mol/µm^2^s. The cells are initially located in a rhombus shape with an equal side length of 21 µm (Figure 6a).

**Figure 6:**
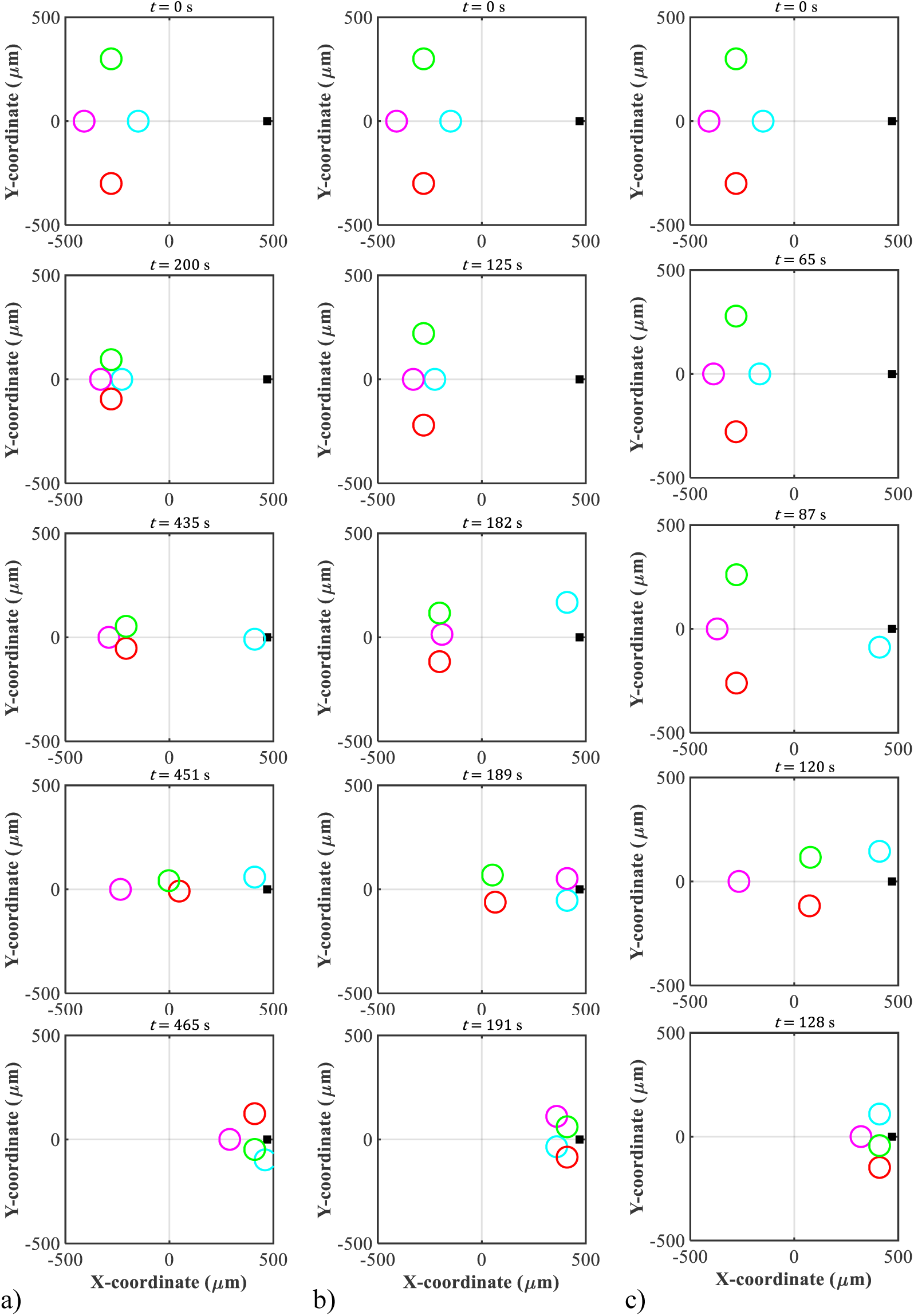
Chemo-mechanically induced collective cell motility for varying growth factor diffusivity. Positions of four cells migrating with fully deterministic motion on a substrate with a constant production rate of α = 8,000 mol/µm^2^s and three different diffusion rates of the growth factor source of *D* = 23 (a), 83 (b), and 179 (c) µm^2^/s. The growth factor source is located in the region X > 0 (right) as a fixed point, i.e., it does not move with time. a) Initial positions of cells at *t* = 0 s. Cells are located in the region X < 0 (left) in a rhombus shape with an equal side length of 327 µm, minor axis length of 260 µm, and major axis length of 600 µm. Cells migrate towards each other (i.e., towards Y = 0) due to the mechanical cues. Cells cluster after *t* = 200 s. and keep moving around each other. Cells move into the growth factor source as a collective at first, then the front cell (i.e., closest cell to the source) moves faster towards the growth factor source and reaches the source at *t* = 435 s (see also supplemental video V6A). Two cluster cells (i.e., closest to the source) move towards the growth factor source, and cells deform (represented as overlap) at *t* = 451 s. Cells keep moving around the growth factor source randomly along the Y-axis. The furthest cell reaches the source last at *t* = 465 s. b) Initial configuration of the cells at *t* = 0 s. Cells migrate towards each other (i.e., towards Y = 0). At *t* = 125 s, neighbouring two cells join at Y = 0. At *t* = 182 s, cells keep moving towards each other in the direction of the growth factor source. The first cell closer to the source reaches the source first, followed by the cell second-closest to the source at *t* = 189 s. Once the cells reach the growth factor source, they move randomly along the Y-direction (see also supplemental video V6B). Finally, the two cells reach the source site last at *t* =191 s. c) Positions of cells at *t* = 0 s. At *t* = 65 s, the cells move towards each other in the direction of resultant strain energy density at first. Once the cells detect the chemical concentration gradient, they move towards the source at *t* = 87 s (see also supplemental video V5C). The first cell closer to the growth factor source reaches the source first, followed by the two cells second closest to the source at *t* = 120 s. At *t* = 128 s, the furthest cell reaches the source last. (Values and sources of the model parameters are provided in Table 1.)

The first case considers the diffusion rate of the growth factor of *D* = 23 µm^2^/s (Figure 6a). The four cells primarily move towards each other. Once the cells reach each other (at Y ≈ 0) after *t* = 200 s, they move collectively in the X-direction towards the growth factor source. Once the concentration gradient of the growth factor starts dominating the mechanical cues, the cell in front detects the concentration gradient first and moves faster than the other three cells towards the growth factor source (*t* = 435 s). The leading cell keeps moving around the source randomly in the Y-direction (see also supplemental video V6A). After that, two of the remaining three cells closest to the source move collectively towards the source at *t* = 451 s. All four cells reach the growth factor source at *t* ≈ 465 s.

For the diffusion rate of the growth of *D* = 83 µm^2^/s, the cells initially move towards each other (Figure 6b). The two neighbouring cells at Y = 0 reach each other first (*t* = 125 s) and start moving towards the growth factor source (at Y = 0) (see also supplemental video V6B). Once the leading cell reaches the growth factor source (*t* = 182 s), the other three cells continue moving towards each other (at Y ≈ 0) due to mechanical cues. The cell closest to the growth factor moves faster than the other two cells and reaches the growth factor source site at *t* = 189 s. Once the cells reach the growth factor source, they move randomly along the Y-direction (see also supplemental video V6B). Finally, all cells reach the source site and interact with each other (represented by overlap) since the repulsive force is not considered (*t* = 191 s).

For *D* = 23 µm^2^/s, the initial most distant cell reaches the growth factor source last (Figure 6a), whereas for *D* = 83 µm^2^/s the two cells with intermediate initial distance from the growth factor source reach it last. This indicates an increasing relative contribution of the mechanical cue to the migration velocity at higher chemical diffusion rates.

For the diffusion rate of *D* = 179 µm^2^/s, the cells initially move towards each other in the direction of the resultant strain energy density gradient, i.e., towards and along Y = 0, respectively (Figure 6c, *t* = 65 s). The cells are influenced by the release of the chemical source before they form a collective, and the concentration gradient of the growth factor starts dominating the mechanical signals (*t* = 87 s). The leading cell reaches the growth factor source first, followed by two of the three cells closest to the source (*t* = 120 s). The most distant cell reaches the growth factor source site last (*t* = 128 s) (see also supplemental video V5C).

### 3.3 Chemo-mechanically induced single-cell motility

#### 3.3.1 Chemo-mechanically induced single-cell motility for varying growth factor productivity

Two cells 539 µm apart on the negative substrate region (X < 0) are considered with three different growth factor production rates of α = 60, 970, and 8,000 mol/µm^2^s and a constant diffusion rate of *D* = 179 µm^2^/s. One cell is immobile, and the other cell is allowed to move. Both cells exert traction forces, and the stationary cell attracts the motile cell due to mechanical cues (Figure 7).

**Figure 7:**
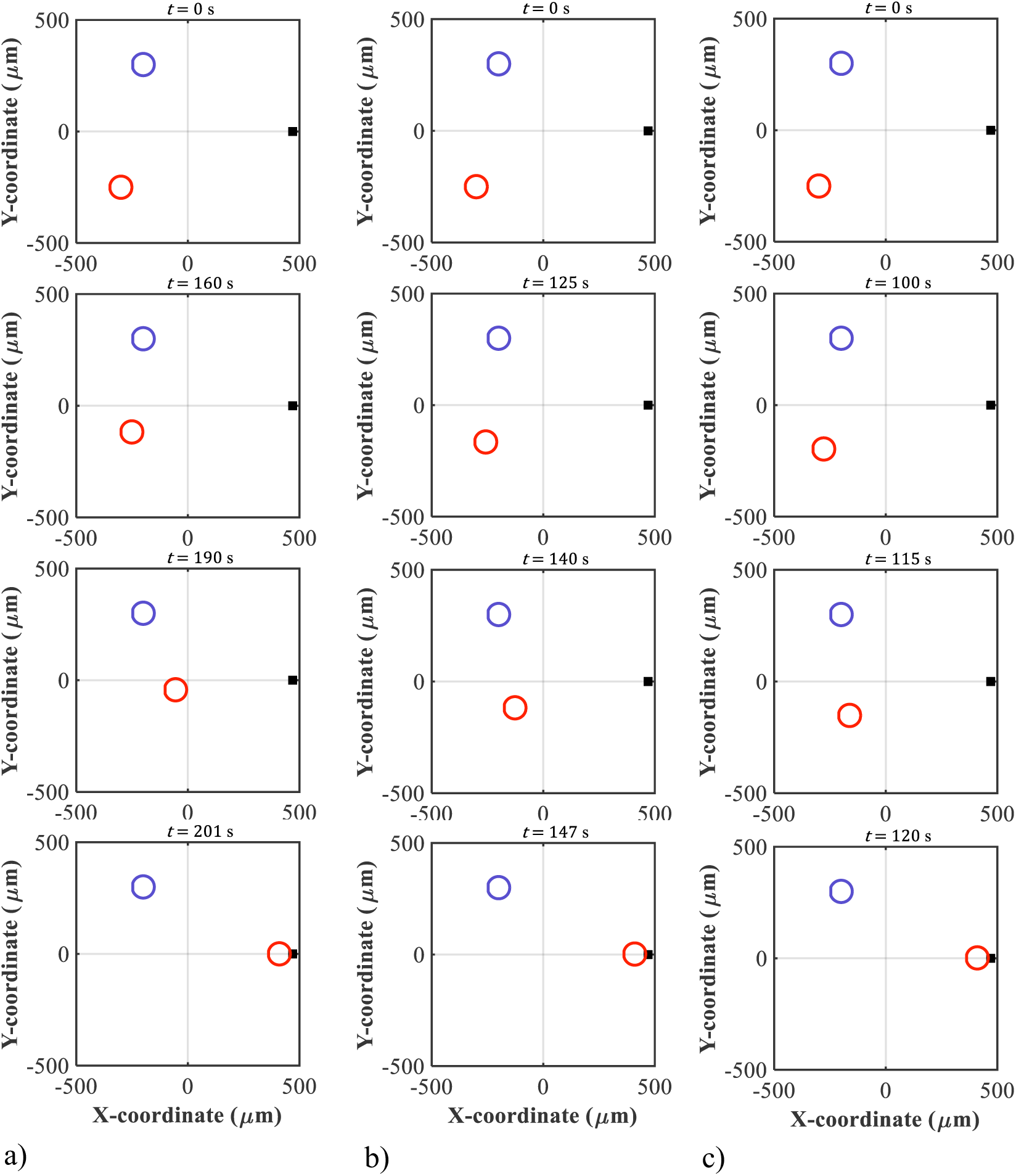
Chemo-mechanically induced single-cell motility for varying growth factor productivity. Positions of a stationary cell (blue circle) and a motile cell (red circle) migrating with fully deterministic motion on a substrate towards a growth factor source (solid square) with a constant diffusion rate *D* = 179 µm^2^/s and different production rates of (a) α = 60 mol/µm^2^s, (b) α = 970 mol/µm^2^s, and (c) α = 8,000 mol/µm^2^s. The source is located at (*x*, *y*) = (470, 0) µm, and cells are initially arranged on the negative side of the X-axis (left). a) Initial position of a motile cell at *t* = 0 s for α = 60 mol/µm^2^s. Firstly, the motile cell moves towards the stationary cell due to mechanical cues at *t* = 160 s. After *t* = 190 s, the cell moves towards the growth factor source. At *t* = 201 s, the cell reaches the source and stays still around it (see also supplemental video V7A). b) Initial position of a motile cell at *t* = 0 s for α = 970 mol/µm^2^s. At *t* = 125 s, the motile cell moves towards the stationary cell (see also supplemental video V7B). After *t* = 140 s, the motile cell moves towards and reaches the growth factor source at *t* = 147 s. c) Initial position of a motile cell at *t* = 0 s for α = 8000 mol/µm^2^s. The cell migrates towards the stationary cell at *t* = 100 s. At *t* = 115 s, the cell moves towards the growth factor source (see also supplemental video V7C). At *t* = 120 s, the cell reaches the source. (Values and sources of the model parameters are provided in Table 1.)

For the growth factor production rate of α = 60 mol/µm^2^s (Figure 7a), the motility of the motile cell (red) is firstly directed towards the stationary cell (blue) due to the initially dominant mechanical cues (*t* = 160 s). After *t* = 190 s, the motile cell moves in the direction of the growth factor source and away from the stationary cell as the growth factor concentration gradient exceeds the mechanical migratory cues (see also supplemental video V7A). The motile cell reaches the growth factor source at *t* = 201 s.

For the growth factor production rates of α = 970 mol/µm^2^s (Figure 7b) and α = 8,000 mol/µm^2^s (Figure 7c), similar cell behaviour is shown as in the previous case. The single motile cell reaches the growth factor source at *t* = 147 s for α = 970 mol/µm^2^s and at *t* = 120 s for α = 8,000 mol/µm^2^s, indicating a higher migration velocity at higher *α*. The cell moves faster towards the growth factor source at higher α followed by a plateau (see also supplemental videos V7B and V7C).

#### 3.3.2 Chemo-mechanically induced single-cell motility for varying growth factor diffusivity

One motile and one stationary cell on the negative region (X < 0) of the substrate, with a distance of 539 µm between the cells, are considered with three growth factor diffusion rates *D* = 23, 83, and 179 µm^2^/s and a constant production rate of α = 8,000 mol/µm^2^s.

For the diffusion rate *D* = 23 µm^2^/s (Figure 8a), the motility of the motile cell is initially towards the stationary cell due to the mechanical cues. The motile cell reaches the stationary cell at *t* = 297 s and keeps moving around the stationary cell (*t* = 424 s) (see also supplemental video V8A). Thereafter, the cell’s movement changes towards the growth factor source as the chemical signal dominates the mechanical cues. After *t* = 436 s, the motile cell reaches the growth factor source.

**Figure 8:**
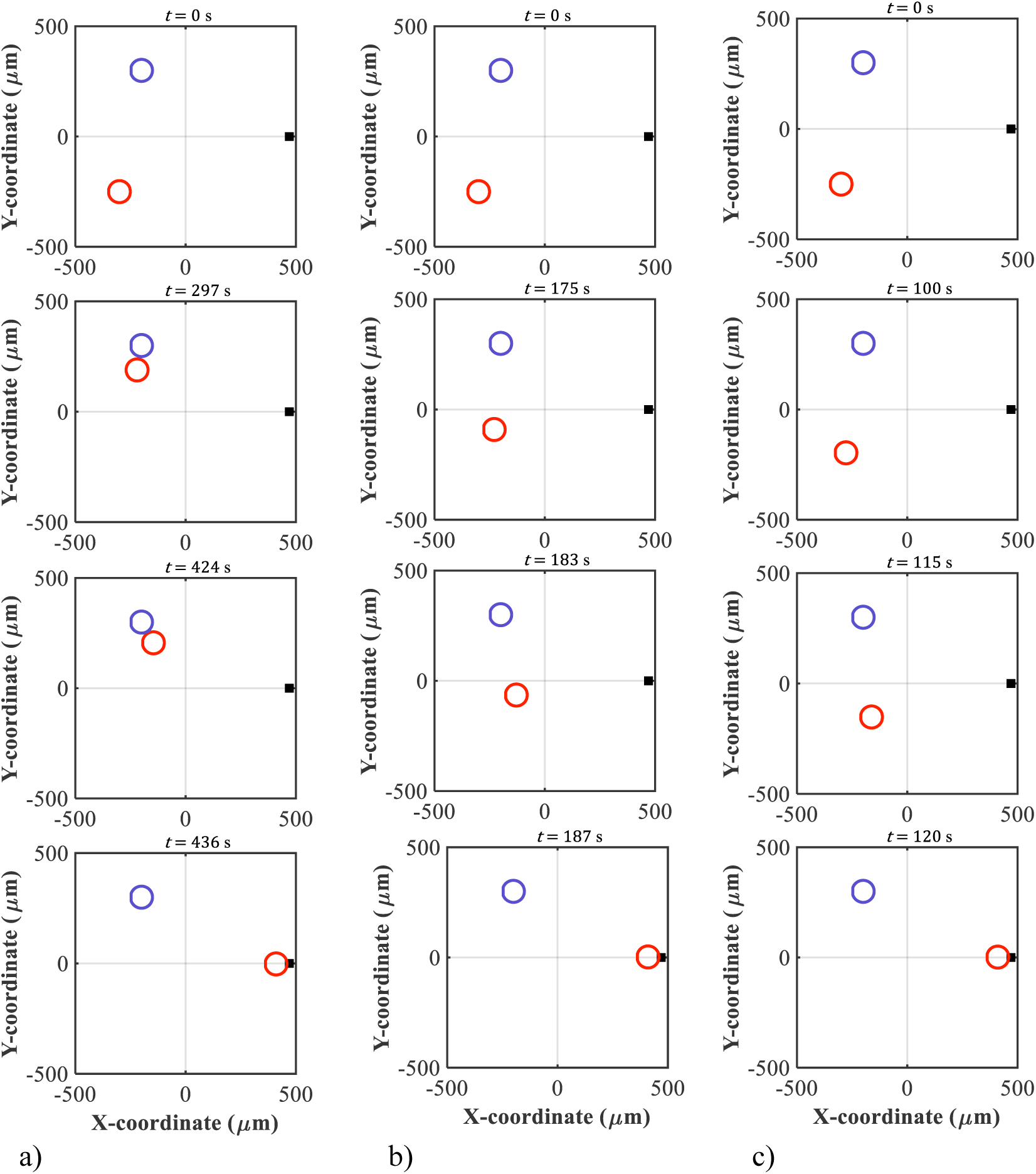
Chemo-mechanically induced single-cell motility for varying growth factor diffusivity. Positions of a stationary cell (blue circle) and a motile cell (red circle) during motility with fully deterministic motion on a substrate towards a growth factor source (solid square) with a constant production rate α = 8,000 mol/µm^2^s and different diffusion rates of (a) *D* = 23 µm^2^/s, (b) *D* = 83 µm^2^/s, (c) *D* = 179 µm^2^/s. The source is located at (*x*, *y*) = (470, 0) µm, and the initial positions of the cells at *t* = 0 s on the substrate region with X < 0 (left) are the same for the three cases. a) For *D* = 23 µm^2^/s, the motile cell migrates towards the stationary cell due to mechanical cues at *t* = 297 s. After *t* = 424 s, the motile cell changes motility towards the growth factor source (see also supplemental video V8A). At *t* = 436 s, the motile cell reaches the growth factor source. b) For *D* = 83 µm^2^/s, the motile cell migrates towards the stationary cell at *t* = 175 s. After *t* = 183 s, the motile cell moves towards the growth factor source and reaches the source at *t* = 187 s (see also supplemental video V8B). c) For *D* = 179 µm^2^/s, the motile cell moves slightly towards the stationary cell at *t* = 100 s. After *t* = 115 s, the motile cell moves towards the growth factor source (see also supplemental video V7C). At *t* = 120 s, the cell reaches the source. (Values and sources of the model parameters are provided in Table 1.)

For *D* = 83 µm^2^/s (Figure 8b), the initial motility of the motile cell towards the stationary cell and the growth factor source due to mechanical and chemical cues is observed around *t* = 175 s. After *t* = 183 s, the motile cell starts moving towards the growth factor source (see also supplemental video V8B). The motile cell reaches the growth factor source at *t* = 187 s and remains still. Compared to *D* = 23 µm^2^/s, the motile cell does not reach the stationary cell, indicating an increasing effect of the chemical cue on cell motility with an increase in the growth factor diffusion rate to *D* = 83 µm^2^/s.

For *D* = 179 µm^2^/s (Figure 8c), the motile cell shows only a minute movement towards the stationary cell (*t* = 100 s), indicating that the growth factor concentration gradient is more dominant than the mechanical cue. Therefore, the cell moves towards the growth factor source (*t* = 115 s) and reaches the source at *t* = 120 s (see also supplemental video V7C).

### 3.4 Quantitative validation

The calibrated model predicts that the average migration velocity depends on the growth factor production rate and diffusion rate (Figure 9). Importantly, the calibrated velocities lie within the reported experimentally determined velocity range for fibroblasts [50, 51], demonstrating quantitative agreement in magnitude and parameter dependence.

**Figure 9:**
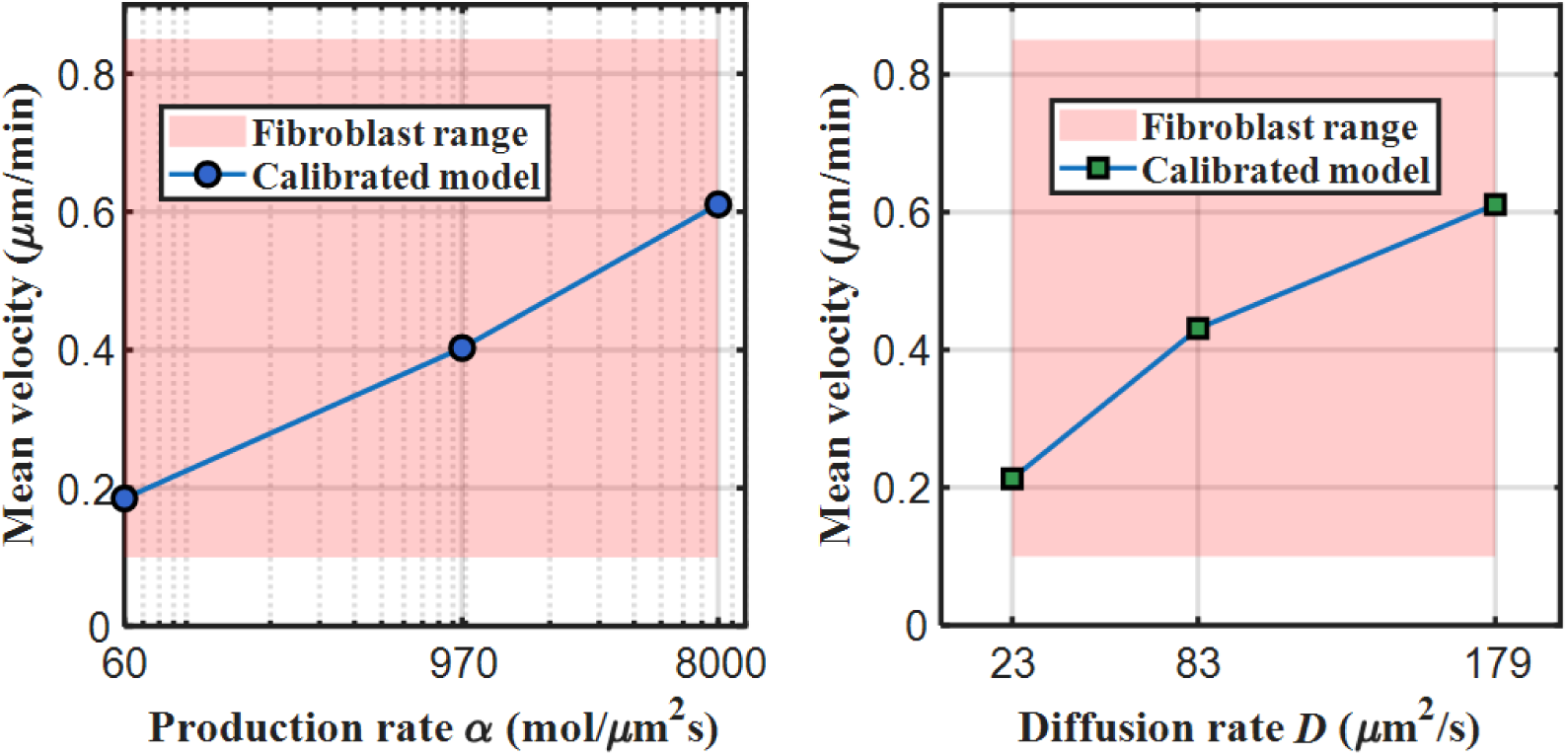
Comparison between calibrated model (calibrated mobility parameter σ_cal_ = 2.76 µm^4^/s mol) predictions and experimental data for the migration velocity of fibroblasts. (a) Mean migration velocity versus growth factor production rate of α = 60, 970, and 8000 mol/µm^2^s and a diffusion rate *D* = 179 µm^2^/s. (b) Mean migration velocity versus growth factor diffusion rate of *D* = 23, 83, and 179 µm^2^/s and a production rate *α* = 8000 mol/µm^2^s. The shaded regions indicate the experimental range of fibroblast migration velocity (≈ 0.1-0.85 µm/min [50, 51]). After calibrating the mobility rate parameter σ_cal_, the predicted mean velocities fall within the experimental range for the production and diffusion rates of the growth factor, demonstrating quantitative agreement in magnitude and parameter dependence. (Values and sources of the model parameters are provided in Table 1.)

## 4 Discussion

The developed model describes the mechanically induced cellular motion on a two-dimensional (2D) planar substrate with a growth factor gradient. The formalism is different from those proposed previously [15, 31, 32, 34, 48, 56] as growth factor productivity and diffusivity are considered to be parameters determining cellular motility. Therefore, the proposed model allows studying how the evolution of chemical concentration gradients guides cell motility during mechanical cellular interactions.

The growth factor concentration gradient is assumed to determine cellular motion in a discrete time. Therefore, diffusion of the growth factor in the substrate is modelled by the transient diffusion of Green’s function on a continuum. The diffusion of the growth factor is monitored using the fundamental solution of the Green function, which determines the concentrations only at the cells’ positions, without the requirement of solving the entire computational field using other discretisation approaches, such as the finite element approach [57–60]. Using the fundamental solution of Green’s function requires elaborate time integration techniques to (i) estimate Green’s function and (ii) track the computations of the cells’ paths towards the growth factor source. However, this approach improves the numerical efficiency under qualitative precision levels by simplifying the diffusion process for semi-bounded and infinite domains compared to approaches that require solving the entire diffusion field using discretisation techniques.

Boundary conditions, such as zero flux diffusion or recurrent conditions, are essential to represent physical situations for finite bounded domains. The mathematical requirement to formulate boundary conditions is often not desirable when describing biological processes for larger diffusion fields [61]. However, the fundamental solution of Green’s function enables the evaluation of the growth factor concentration at any point of the substrate and at any time. Similar numerical approaches have been used to model cancer cell motility [42, 47, 62] and wound closure [43, 63].

Several *in vitro* studies on collective cell motility in the presence of chemical sources demonstrated that cell motility is influenced by the concentration of chemical sources and the strength of chemotactic signals [64, 65]. Nevertheless, the role of the productivity and diffusivity of such chemical sources remains poorly understood and requires further investigation. Even though the formalism of the proposed model uses a simple approach, multiple biological processes can be easily incorporated into the developed model.

For the motility of a single cell towards a growth factor source, the proposed model predicted a decrease in the motility time for an increase in the growth factor productivity and diffusivity (Figure 2a and b). This prediction agrees with experimental observations of a decreasing motility time with increasing chemical signal strength for several cell types [64, 66, 67]. The decreasing motility time correlates with an increase in the average motility velocity of a single cell, i.e., from the start of growth factor release to the arrival of the cell at the growth factor source, for increasing growth factor productivity and diffusivity (Figure 2c and d). These findings are qualitatively similar to experimental results for increasing chemical signal strength [65, 67, 68]. However, the decreasing maximum motility velocity of a single cell with increasing growth factor diffusivity (Figure 2d) contrasts with the decrease in motility duration. This observation may be explained by the inverse relationship between the diffusion coefficient and concentration gradient, i.e. an increase in diffusivity leads to a decrease in the concentration gradient and, hence, a weaker chemotactic cue [38, 64]. It is worth noting that neither higher productivity nor lower diffusivity necessarily leads to an increase in the maximum motility velocity. This behaviour may be explained by the dependence of cell velocity on the effective slope of the growth factor gradient, which increases with growth factor signal strength but eventually reaches a plateau due to limitations in gradient sensing [69]. Therefore, further increases in signal strength do not translate into higher migration velocities. This mechanism may explain the model prediction, at least in part.

For collective cell motility in the presence of a growth factor but in the absence of mechanotactic cues, the developed model predicts an increase in motility velocity with increasing growth factor productivity and diffusivity. Such motility behaviour has been reported in experimental [10, 11, 15] and numerical studies [21, 32, 49]. Cells in a collective only start migrating once they sense the growth factor gradient. The cells closest to the growth factor source start moving towards the growth factor source first, with start time and motility velocity depending on the growth factor productivity (Figure 3) and diffusivity (Figure 4). The cell motility is affected more by a change in diffusivity than in productivity of the growth factor: The motility duration decreases approximately two-fold for a 2.2-fold increase in growth factor diffusivity (Figure 4b and c) compared to a 130-fold increase in growth factor productivity (Figure 3a and c). The growth factor diffusivity determines the propagation velocity of the growth factor molecules in the substrate. For an increase in diffusivity, the growth factor molecules propagate faster; the growth factor gradient reaches or can be sensed by the cells earlier, and cell motility starts earlier.

The model results reveal several differences between cell motility behaviour due to combined chemical and mechanical cues and motility induced by chemical cues only. The chemo-mechanical motility of multiple cells is directed towards the growth factor source but may be distinguished into two phases relative to the growth factor release. Initially, the cells migrate towards each other due to the dominance of the mechanical cues (exerted by the cells) over the chemical cues from the growth factor source (Figure 5 and Figure 6). In this phase, the motility is predominantly mechanically induced. As more growth factor is released from the source and diffuses in the substrate, the strength of the growth factor gradient increases and the chemical cue may dominate the mechanotactic cue. At this stage, the cells start migrating towards and eventually reach the growth factor source. Similar behaviour is reported in experiments [16, 70–72].

Comparing the chemo-mechanical motility of one motile cell in the presence of one stationary cell (Figure 7) to that of multiple motile cells (Figure 5) indicates that the mechanical interactions amongst multiple motile cells delay their arrival at the growth factor source, i.e. result in slower collective cell motility compared to the single cell motility. Even though the predicted dependence of cell motility on mechanotactic cell interactions can be ascribed to the initial cell positions [8], these interactions do not affect the final stage of the motility, i.e., the arrival of the cells at the growth factor source. These results agree with experimental [73, 74] and numerical studies [21, 30, 31, 75].

As observed for the chemically induced cell motility without mechanical cues, chemo-mechanical cell motility is also more affected by the variation of the growth factor diffusivity than the growth factor productivity. The motility duration decreases approximately 1.5-fold for a 2.2-fold increase in growth factor diffusivity (Figure 6b and c) compared to a 130-fold increase in growth factor productivity (Figure 5a and c).

The present model predicts migration velocity as a function of the growth factor gradient and the distance between the cell and the growth factor source. As a result, the model overestimates migration velocity relative to experimentally observed fibroblast motion on planar substrates. This behaviour reflects the absence of an explicit saturation mechanism in the current formulation, in which velocity increases linearly with the chemotactic driving force.

The quantitative validation indicates that the model captures the essential physical and biological mechanisms governing chemo-mechanically induced cell motility, including sensitivity to gradient strength and transport properties. Before calibration, the predicted velocity range of 8-30 µm/min was closer to velocities observed in fast-migrating cell types such as keratocytes (≈ 5-20 µm/min [52–54]), which highlights the role of scaling in reconciling model outputs with cell type-specific motility. Overall, the calibrated model provides a consistent description of fibroblast-type motility while preserving its mechanistic basis. Despite this difference in magnitude, the model produces experimentally observed trends, including increased migration velocity with increasing growth factor stimulation, as observed experimentally for fibroblast migration [55]. These findings indicate that the model captures the underlying physical and biological mechanisms governing chemo-mechanically induced cell motility, while requiring calibration for the representation of fibroblast migration speed.

Although the results generated with the proposed model are in agreement with previously reported experimental data for cell motility in complex environments, the model has some limitations. The model approach does not consider the maximum migration rate towards the growth factor source. The signal strength that attracts the cells increases over time since the growth factor source is immobilised and non-soluble. In contrast, experimental studies reported that cells have limited velocity to move towards the source [10, 14, 71].

Future extensions of the model may incorporate a saturation mechanism in the chemotactic response, for example, due to receptor saturation or bounded sensitivity to the growth factor gradient, to limit the migration velocity under strong stimuli.

The model uses a simple approach to describe the cell motion once the cells reach the growth factor source. The signal strength that attracts the cell becomes very large since the growth factor source is immobilised and non-soluble. In addition, the distance between the cells and the growth factor source, described in Eq. (6), becomes very small.

The model formalism considers the growth factor productivity and diffusivity to be constant over time during the migration process. In contrast, *in vitro* studies reported that these parameters depend on the composition of the growth factor [31, 64] and the substrate [67, 76]. Therefore, considering the productivity and diffusivity of the growth factor as linear and isotropic is a simplification. Implementing anisotropic growth factor productivity and diffusivity may need a different numerical approach or an additional numerical scheme in the current approach in future studies.

The model includes some assumptions to simplify the approach, such as a circular cell shape, whereas the cell shape changes during motility [77]. Also, the number of cells for the chemo-mechanical simulations is limited to four, and cell deformation has been incorporated as partial overlapping. Moreover, stochastic processes, such as randomness of cell motion and cell proliferation, death, and growth, have not been considered in the current model. However, the model performs well as it captures the essential features, such as the mechanotactic cell-cell interactions and chemically induced cell motility.

Even though the chemical interactions between cells have not been considered, an extension of the model will not change the findings. However, the complexity of the model will increase, and more parameters will need to be determined from experiments.

## 5 Conclusion

A mathematical model for cell motility has been developed based on two stimulus types, i.e., chemical and mechanical. For a weak chemical concentration gradient, the cells migrate towards one another due to the inter-cellular mechanical cues. In contrast, cells migrate towards the chemical source for a strong chemical concentration gradient that dominates the mechanical signals. This model offers a tool to predict collective cell motility *in vivo* or *in vitro* due to the flexible approach to consider the effects of several cues on cell motility simultaneously.

## Funding

The research reported in this publication was supported financially by the Organization for Women in Science for the Developing World and Swedish International Development Cooperation Agency (doctoral scholarship to RA), the European Mathematical Society (collaborative research visit award to RA), the South African Medical Research Council (grant SIR328148 to TF), and the National Research Foundation of South Africa (grants UID 92531 and 93542 to TF). Any opinion, findings, conclusions and recommendations expressed in this publication are those of the authors and therefore the funders do not accept any liability in regard thereto.

## Conflict of Interests

Conflicts of interest do not exist.

## Author Contributions

RKA: Conceptualization, Data curation, Formal analysis, Funding acquisition, Investigation, Methodology, Project administration, Software, Validation, Visualization, Writing – Original Draft, and Writing – Review & Editing.

TA: Conceptualization, Methodology, Project administration, Supervision, and Writing – Review & Editing.

NHD: Conceptualization, Methodology, Supervision, and Writing – Review & Editing.

FV: Conceptualization, Methodology, Writing – Review & Editing.

TF: Conceptualization, Funding acquisition, Methodology, Project administration, Resources, Supervision, Validation, Visualization, and Writing – Review & Editing.

## Data availability

Software code, data, and videos supporting this article are available on the University of Cape Town’s institutional data repository (ZivaHub) at http://doi.org/10.25375/uct.25507387 as Ahmed RK, Abdalrahman T, Davies NH, Vermolen F, Franz T. Software code, data, and videos for A Mathematical Model for Chemo-mechanically Induced Collective Cell Motility on Planar Elastic Substrates. ZivaHub, 2024, DOI: 10.25375/uct.25507387.

## Ethics statement

None

## Notes

http://doi.org/10.25375/uct.25507387

### Competing Interest Statement

The authors have declared no competing interest.

### Summary of Updates

Minor revision of a statement in results section 3.1.1 in the paragraph preceding Figure 2 and in the caption of Figure 2.

